# Community engagement through “student-led science” for dengue prevention during the COVID-19 pandemic in Córdoba, Argentina

**DOI:** 10.1101/2022.10.04.510837

**Authors:** Elizabet L. Estallo, Magali Madelon, Elisabet M. Benitez, Anna M. Stewart-Ibarra, Francisco F. Ludueña-Almeida

## Abstract

**Background:** During 2019-2020 while COVID-19 pandemic, the Americas were facing the biggest dengue fever epidemic in recent history. Traditional vector control programs, based on insecticide application have been insufficient to control the spread of dengue fever. Several studies suggest refocusing on education with the aim of an integrated vector management strategy within the local ecological-community context. We aim to assess community perceptions, knowledge, attitude, preventive practice, and action through student-led science assignments regarding dengue fever, prevention, and socio-ecological factors in temperate Córdoba, Argentina.

**Methods:** The study was conducted during the COVID-19 quarantine when schools switched to online education for the first time. Several activities through Google Classroom platform included a survey to one student’s family member, and an outdoor activity to assess their attitudes and to clean the backyard and gardens.

**Results:** Significant number of respondents developed good preventive practices and increased their knowledge about the vector and disease highlighting that 75% of responders knew that dengue fever was transmitted by a mosquito, 81.96% declared having obtained knowledge regarding dengue and vector through television, 56% affirm that dengue is a severe illness, 67% of respondents admitted that individuals play an important role in the prevention of dengue. Regarding mosquito control activities, 90% of respondents reported turning containers.

**Conclusions:** This highlights the need for school programs with curricula to address vector biology and the prevention of vector-borne diseases not only during activity periods when mosquitoes batter people but all year long to do real prevention.

## Introduction

Since 2020 the COVID-19 pandemic has affected countries around the world. Argentina has been in mandatory quarantine since March 20, 2020, being extended several times in the year [1] with online school and the emergence of online education and opportunities and challenges faced. At the same time, the Americas were also facing the biggest dengue fever (DEN) epidemic in history [2]. Dengue fever is caused by dengue virus (DENV), transmitted by *Aedes aegypti* mosquito to humans, with a wide geographic spread of the virus and its vector in tropical, subtropical, and even temperate areas [3, 4]. During the season 2019-2020 Argentina has reported 58,889 DEN cases, with 3 circulating serotypes in the country (1, 2, 4) and 26 deaths. This season exceeded by almost 40.5% accumulated cases in the 2015-2016 season, being the largest dengue outbreak that has been recorded in the history of the country so far [5]. It should be noted that the temperate central region of Argentina has been one of the most affected, with almost 40% of the cases reported. In Córdoba province, located in the temperate central area, 3,631 DEN cases were reported with two circulating serotypes (1 and 4) [3, 5]. Unlike the 2019-2020 season, during the last season, 2020-2021 were reported 4,653 DEN cases (serotypes 1, 2, 4) in all the country [6]. Several challenges must be sorted by the local health system, such as the focal control, because of the preventive isolation, as well as the difficulty of the clinic diagnostic by the physicians due to similar febrile symptoms (Federico Layun, from Cordoba municipality epidemiological system, personal communication).

Rapid human development, changes in demographics, population, rural-urban migration, inadequate basic urban infrastructure, an increase of solid waste, such as discarded plastic containers and other abandoned items such as cars wheels, plastic bags, unused objects that can provide larval mosquito habitats, and become a risk factor when DENV is circulating. These risk factors combined with appropriate environmental conditions enhance the likelihood of viral transmission [7, 8].

Traditional vector programs, based on chemical intervention, are not enough [9], therefore several studies suggest focusing on education with aim of an integrated management strategy (IMS) within the ecological-community context. In temperate areas like Cordoba, education is focused just during the high vector activity and not all year long as could be observed in some tropical countries. The IMS seeks to modify the behavior of individuals and the community in such a way as to reduce risk factors for transmission with coordinated measures both within and outside the health sector [10]. Indeed, there is evidence that education can lead to behavior changes related to DEN, revealing that knowledge scores were significantly increased after health education programs, barring a language communication gap [11, 12]. Prior studies have involved student’s community and their families, focusing on dengue prevention, health promotion, and action, as in Honduras, where educational interventions were applied as part of a comprehensive plan for the control of *Ae. aegypti*, allowing the teachers and families in the program, inducing their participation in reducing sources at home [13]. In Mexico through the implementation of an educational strategy, knowledge, attitudes, and practices about self-care in their schools engaged students as dengue promoters’ changers in their homes [14]. The effectiveness of educational interventions was also evaluated in Colombia, where a high percentage of the participants change behavior related to vector breeding places, washing, and covering water storage tanks, containers, as well as collecting unusable potential breeding places in their homes [15]. In Argentina, research in Buenos Aires conducted surveys, interviews, home search for immature mosquitoes, placement of ovitraps (manufactured by students), workshop series, and illustrative classes with teachers and students involving their families and neighbors [16].

Community engagement, through “student-led science”, is a novelty in the area, and could fill the gap on getting to the households, leading to a change in attitudes in the home due to students becoming good household health educators. This study aims to assess household perceptions, knowledge, attitude, preventive practice, and action through student assignments regarding dengue fever, prevention, and socio-ecological factors during the concurrent dengue and COVID-19 pandemic in the city of Cordoba, Argentina, a temperate region where dengue has emerged over the last decade.

## Methodology

### Study area

The city of Córdoba is in the central area of Argentina with the Suquía River crossing it from northwest to east (Fig. 1). It has an area of 576 km2 with a population of 1.3 million people, being the second-largest city in the country, following Buenos Aires. Córdoba is in a temperate region that experiences dry winters, annual rainfall of 800 mm, and an average annual temperature of 18 °C(maximum of 27 °C and minimum of 11 °C) [17].

**Fig. 1.**
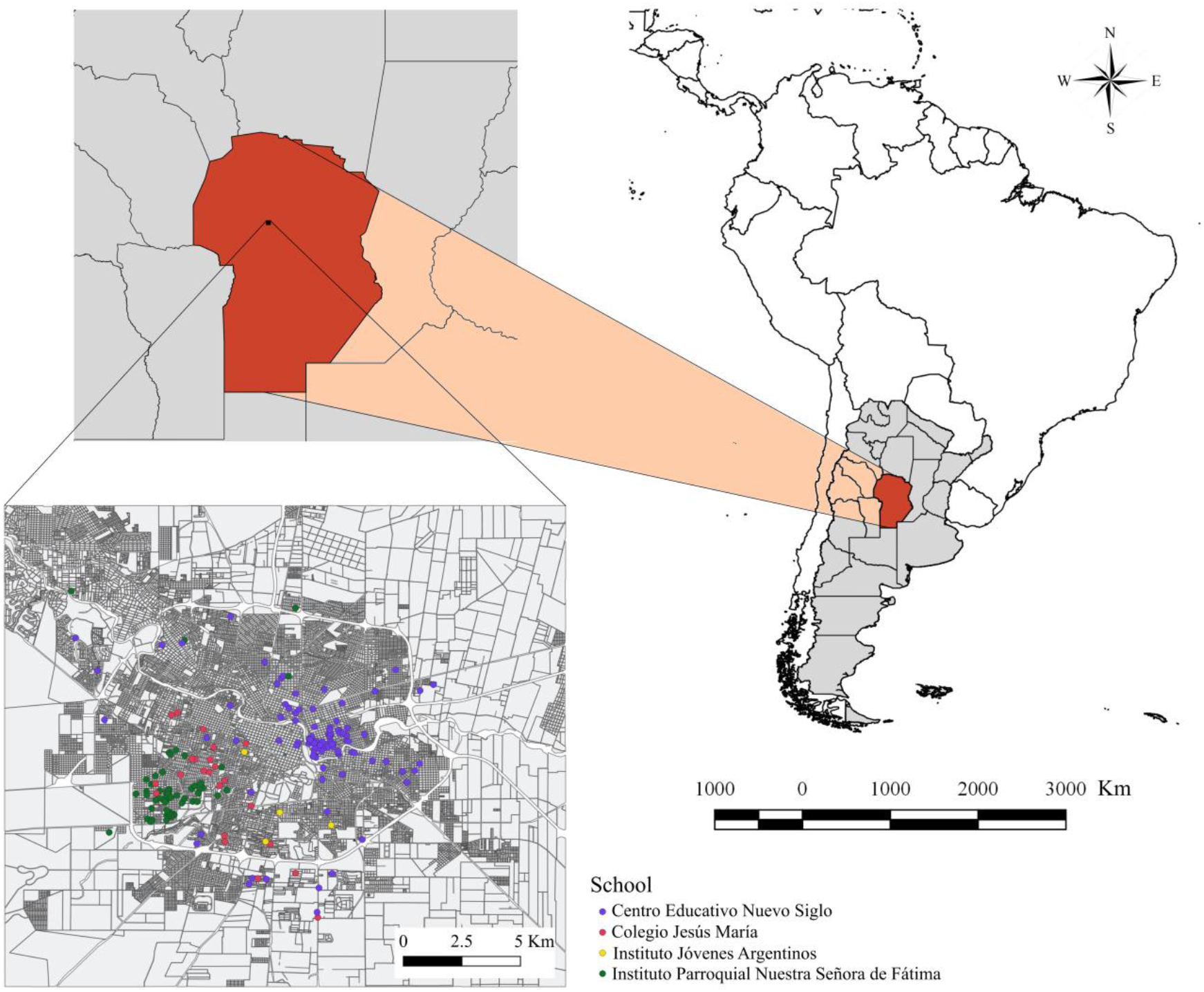
Study area. Location of the Córdoba city (Argentina) where the surveys were conducted during 2020. The points show the approximate location of the respondents and the colors indicate the school to belong the students who conducted the surveys.

Dengue fever emerged in Córdoba over the last decade, with the first dengue outbreak occurring in 2009 [18]. The emergence of dengue has raised concern due to the rising public health burden [3, 19]. Vector control is conducted by the National Health Minister and consists mostly of focal control around homes with dengue cases, including indoor and outdoor fumigation, larviciding with Bacillus thuringiensis israelensis (BTI), and eliminating standing water [20]. In general terms, there have been public education campaigns to prevent dengue transmission; however, in some cases, campaigns have resulted in greater confusion by naming the mosquito as the disease and not differentiating between the virus and vector. In 2014 the government of Cordoba issued a strategic plan called “Plan estratégico de abordaje integral para la prevención y el control del dengue y de la chikungunya en Córdoba” (Strategic plan for integral management for the prevention and control of dengue and chikungunya in Córdoba) [21] that included in the need to strengthen education on dengue and chikungunya in the elementary, middle and high schools, as well as universities and teacher career training (in objective 5 section 6.5.2). Today there are no formal education programs that involve the students and the community and aim to cause a behavioral change to reduce disease risk. Some high schools with specialized Natural Science educational orientations do include in the 4th year of high school (students around 15 years old) some information about the transmission and prevention of relevant diseases in the country, like chagas, hemorrhagic fever, dengue, among others. In February 2020 the provincial government through the memorandum 02/2020 [22] informed school principals of the need for activities related to the dengue epidemic ongoing in the province, where they mention the school as the place to do prevention against dengue and advise to use pedagogical materials that were made available by the education minister of the Cordoba province government.

### Ethics

This study was approved by the ethical committee from National University of Córdoba, Hospital Nacional de Clínicas (Clínicas National Hospital), General coordinator Dra. Susana del Carmen Vanoni. This study was an educational intervention implemented in schools and developed with teachers as part of the curriculum with informed consent signed by each of the school’s principals involved in this research; Lic. Natalia Magalí Carbo (Centro Educativo Nuevo Siglo), Lic. María Inés Buffa (Instituto Jóvenes Argentinos), Prof. Lic. Carlos E. Carranza (Instituto Parroquial Nuestra Señora de Fátima) and Elsa Ferrer (Instituto Jesús María). The study complies with all regulations and confirmation that informed consent was obtained from the participants.

### Field data collection

Because of the mandatory quarantine during the COVID-19 pandemic, the program for developing community engagement by the schools through “student-led science” had three main activities that were labeled as “Dale Block al Dengue” (Spanish) that in English loosely translates to “Let’s Block Dengue,” playing with the social media meaning of blocking a person when you don’t like what they post and translating this to the role of mosquito as dengue vector, knowing that we can block dengue circulation if we eliminate the breeding places for mosquitos to develop. The student science, therefore, consisted of:

1-Survey about perceptions and knowledge regarding dengue fever, prevention, and socio-ecological factors; 2-Activity in their backyards to assess attitude; 3-The activity involving the action of cleaning backyard and gardens to remove unused artificial containers that were potential mosquito habitat. The three activities were developed using the Google Classroom platform. These activities were developed in cooperation and coordination between researchers and high school teachers to assess student science engagement. We worked with four private schools in the city (Fig. 1). One of the school principals decided to involve all classes, therefore every student had to be involved and do the activity as part of the school cooperation and engagement. Each of the other three schools involved only one class related to ecology, environment, and social development science for the natural science orientation high schools. This methodology was developed considering the ongoing dengue outbreak situation in the city and the concomitant mandatory quarantine faced in the country.

### The survey

The survey was generated by Google Forms and put inside the Google Classroom platform to be implemented by the students of the four different private high schools involved in the program. According to the protocol established, each student surveyed a family member >18 years of age, living together in the same home, and one person was surveyed per household. The instructions included not to look in a book or the internet for the answers as it is important to know what is really known about the subject by the respondents. Based on prior field experience in the city and years of collaboration with the province of Córdoba Ministry of the Health (MOH) on the dengue program [23], we determined which variables to include in the survey for community engagement (Additional file 1). **Socio-ecological factors** covered the socio-demographic characteristics consisting of age, highest education degree, relationship with the student doing the survey, self-reported prior dengue infections in household members or knowledge of any person with the infection, people living in the household, travel history to neighboring countries or any other with the vector-borne transmission.

Questions focused on **Vector-borne diseases knowledge** included whether they were aware of dengue, symptoms, and *Ae. aegypti* vector knowledge. We also asked questions regarding their **perception of dengue risk**, the perception of prevention responsibility, and how they carry out the **household control activities**. We asked questions about the **environment surrounding the household** to have better information about proximity to potential vector breeding places within 500 meters from the surveyed homes. These include garbage dumps, depots, water channels, and the Suquía River [8], also we asked people where they have seen mosquitoes in their homes. A radius of 500 meters was selected because several studies have shown that females *Ae. aegypti* mosquitoes generally fly 100-500m [24, 25]. Any man-made discarded container in that range, such as abandoned items in urban public areas, could be a potential breeding place for the vector. For each category of the survey, we generated hypotheses to test (Table 1).

**Table 1.** The hypothesis to test the different survey sections developed during the student-let-science study.

| Survey section | Hypothesis to test |
| --- | --- |
| Socio-ecological factor | - Large percentage of surveyed know of dengue cases in their neighborhoods.<br>- Most trips to risk areas are made in summer vacations. |
| Vector-borne diseases knowledge | -In the population, the correct knowledge of the symptoms of dengue disease depends on the educational level reached by each person.<br>- The source of information about *Ae. aegypti* and the diseases it transmits are electronic means rather than formal educational ones. |
| Perception of dengue, household vector prevention and control practices | - Dengue is perceived as a moderate risk disease.<br>- Dengue prevention is a simple task and carrying it out depends on each individual.<br>- The activities related to eliminating containers that are potential breeding sites are the most frequent practices<br>-The use of repellents is not frequent to avoid the presence of mosquitoes. |
| Household environmental surrounding | -knowing the conditions of the surrounding environment could help with prevention attitudes.<br>-knowing proximity to abandoned areas or rivers could lead to taking preventive actions. |

### Attitude Activity

The second activity involved the students’ attitude of exploring their backyards, gardens and inside their homes to search for containers that hold water and could be potential mosquitoes breeding places. The students received training through lectures and a mosquitoes workshop showing them eggs, larval, pupa, and adult conversed stages to observe in the class. All this training was done during 2019, with the same schools and teachers involved in a citizen science project with our research group. Although, students were trained and taught again online during quarantine about mosquito *Ae. aegypti* life cycle and potential breeding places (Additional file 2). Each one of the students took photographs of their backyards/gardens showing if there were or were not potential mosquito breeding sites. Also, they take photographs of each one of the containers with or without water as well as the mosquito’s stages inside of the water containers. We assess their attitude with the activity done and the photographs sent to the teacher class involved.

### Getting in the action

The last activity involved having the students remove found containers with water that held any arthropod or could be a potential place for mosquitoes to lay eggs becoming a breeding place around their homes. This activity was done once for the school activity requirement, following the survey done before. The students send photographs of each found container, as well as the number of containers in their homes, discriminating between containers with water and without water.

### Analysis

We analyzed each section of the survey using descriptive statistics to show the percentage of responses according to each section above described and we tested several hypotheses for each group of survey sections (Table 1).

Survey responses were coded and grouped as variables: Adequate responses about symptoms (ARS), Adequate responses about dengue and vector knowledge (ARK), and Educational level (EL). For Adequate responses about symptoms (ARS) in relation to Educational level (EL), the ARS was worth 1 point when the respondent answered correctly and 0 points in another case. We tested for the association between ARS and the EL of the respondent using logistic regression analysis. For adequate responses about dengue and vector knowledge (ARK), analyzing the question about how dengue is spread, a score was applied to the answers obtained, attributing a score of 2 points to the answers that included as a response: ^..^through a mosquito bite^..^, or the response ^..^that mosquito should be previously infected^..^, and a score of 5 was assigned to the respondents who answered included the scientific name of the mosquito that transmits dengue, and a value of 10 points when they clarified that ^..^it is the female that bites^..^.

In the same way, a differentiated score was also applied according to the response obtained from the respondents about what *Ae. aegypti* is. We gave a 2 point score to those answers that only mentioned that it was a mosquito or a vector, or that it transmitted Dengue. We gave a 4-point score when they mentioned that *Ae. aegypti* is the scientific name of the mosquito, which transmitted other diseases in addition to Dengue, or specified the physical characteristics of the species.

Finally, a sum of the obtained scores was made, and those respondents who had obtained between 10 and 28 points were considered as answers with sufficient knowledge. And those who had obtained less than 10 points as answers with insufficient knowledge. These results were analyzed according to the age of the respondent and the educational level reported.

Preventive practices refer to the interviewee’s self-reported actions to avoid breeding mosquitoes and mosquito bites in their dwellings. Among actions declared were mowing the grass, the use of mosquito’s coils as repellents and spirals, fumigation, physical elimination of habitats to reduce breeding sites, and chemical products applied in swimming pools. According to the expert consensus regarding *Ae. aegypti* control [26], not all the mentioned actions have the same weight. Therefore, we calculated a weighted score of control actions: *V* = 10*R* + 5*B* + 5*P* + 2*F* + *S*

Where *V* is the total points assigned to the surveyor according to the declared actions, represented with letters (*R, B, P, F, S*) receiving 1 point if the action was declared in the answer as done, or 0 points if the action wasn’t declared.

The *R* indicated the action against containers (physical elimination of habitats, cover containers, put down artificial containers to reduce breeding sites) which was scored with 10 points because is the most action to reduce *Ae. aegypti* population. *B* indicated bite protection against mosquitoes through the use of insecticides like repellents and spirals, and this action was valued with 5 points. *P* corresponds to applied chemicals in swimming pools that were valued with 5 points. *F* refers to fumigation or spraying the dwelling and peri-dwelling with insecticides, which was valued with 2 points. Even though with this practice it is possible to eliminate mosquitoes, other insects are affected as well as human health and pets. *S* is assigned to grass mowing, which gets 1 point because the only impact is a reduction of a possible refuge for male or female mosquitoes.

Therefore, the *V* value could be from 0 to 23, and the actions were qualified as sufficient if they scored equal to or greater than 12 points, and insufficient if less than 12 preventive practices were associated with age and education level through logistic regression analysis. All statistical analyses were conducted in Infostat software [27].

## Results

A total of 240 students conducted the surveys, but only 188 accomplish the condition of surveying one family member >18 years of age, living together in the same home. Therefore 188 surveyors were considered in this study, and 52 participants were excluded because either they did not live together in the same home or they were not >18 years of age, with a remaining 188 study participants included in the analysis.

### Socio-ecological factors

Table 2 provides a summary of socio-ecological factors including socio-demographic characteristics of the participants. Most participants were 30 years of age or older (88.2%). Most homes had 4 inhabitants. Although the study was conducted amidst the biggest dengue outbreak in history, more than 90% were unaware of dengue cases in their neighborhoods. Most of the responders had completed high school. Among the people that travel outside the country, most traveled during the months of January and February, the southern hemisphere summer vacation period, to neighboring countries (Table 2).

**Table 2.**
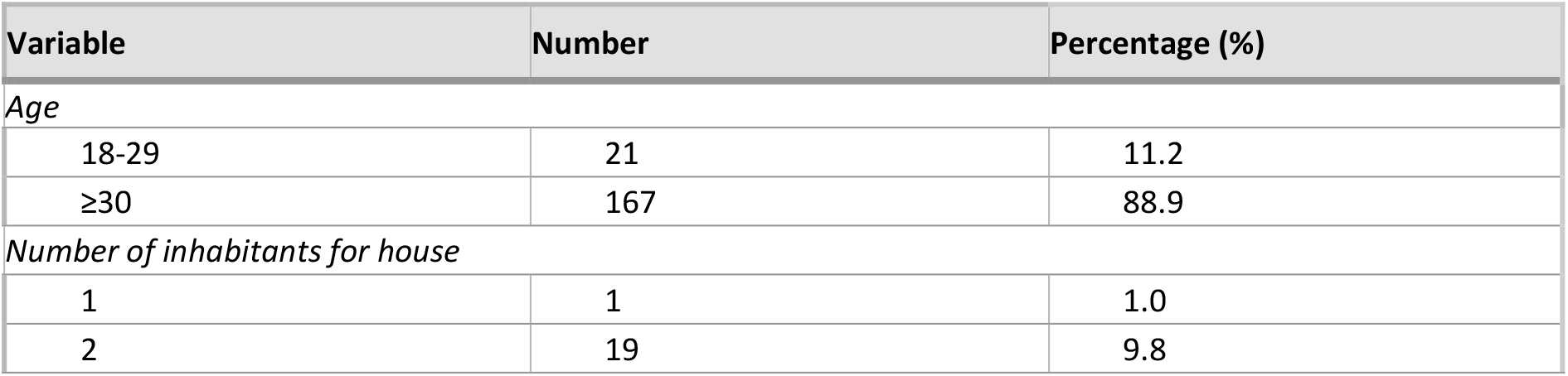
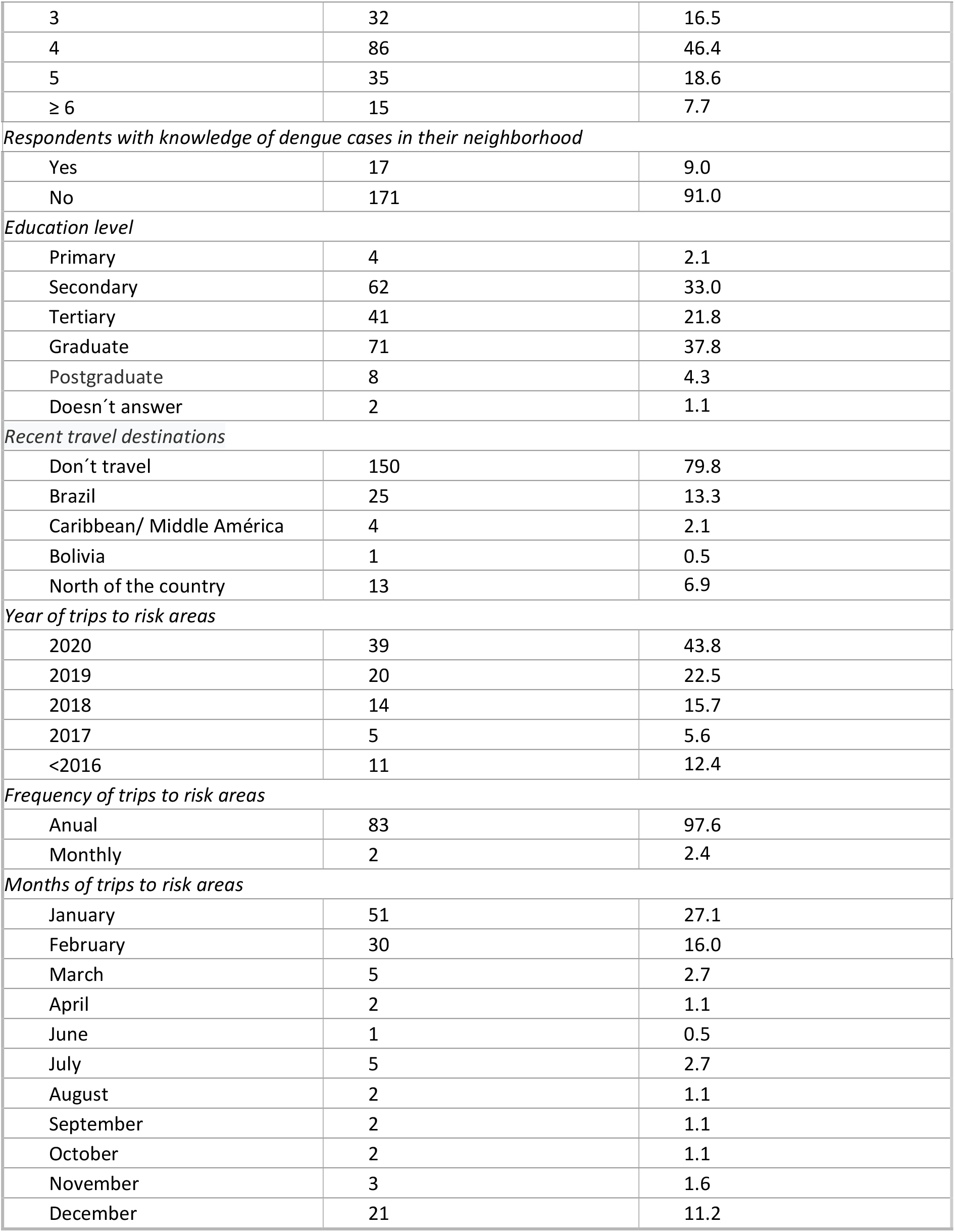
Information provided by respondents on socio-ecological factors.

| Variable | Number | Percentage (%) |
| --- | --- | --- |
| *Age* |  |  |
| 18-29 | 21 | 11.2 |
| ≥30 | 167 | 88.9 |
| *Number of inhabitants for house* |  |  |
| 1 | 1 | 1.0 |
| 2 | 19 | 9.8 |
| 3 | 32 | 16.5 |
| 4 | 86 | 46.4 |
| 5 | 35 | 18.6 |
| ≥ 6 | 15 | 7.7 |
| *Respondents with knowledge of dengue cases in their neighborhood* |  |  |
| Yes | 17 | 9.0 |
| No | 171 | 91.0 |
| *Education level* |  |  |
| Primary | 4 | 2.1 |
| Secondary | 62 | 33.0 |
| Tertiary | 41 | 21.8 |
| Graduate | 71 | 37.8 |
| Postgraduate | 8 | 4.3 |
| Doesn't answer | 2 | 1.1 |
| *Recent travel destinations* |  |  |
| Don't travel | 150 | 79.8 |
| Brazil | 25 | 13.3 |
| Caribbean/ Middle América | 4 | 2.1 |
| Bolivia | 1 | 0.5 |
| North of the country | 13 | 6.9 |
| *Year of trips to risk areas* |  |  |
| 2020 | 39 | 43.8 |
| 2019 | 20 | 22.5 |
| 2018 | 14 | 15.7 |
| 2017 | 5 | 5.6 |
| <2016 | 11 | 12.4 |
| *Frequency of trips to risk areas* |  |  |
| Annual | 83 | 97.6 |
| Monthly | 2 | 2.4 |
| *Months of trips to risk areas* |  |  |
| January | 51 | 27.1 |
| February | 30 | 16.0 |
| March | 5 | 2.7 |
| April | 2 | 1.1 |
| June | 1 | 0.5 |
| July | 5 | 2.7 |
| August | 2 | 1.1 |
| September | 2 | 1.1 |
| October | 2 | 1.1 |
| November | 3 | 1.6 |
| December | 21 | 11.2 |

### Vector-borne diseases knowledge

Most of the responders (75%; 141/188) knew that dengue fever was transmitted by a mosquito. For knowledge on dengue symptoms, fever, body ache, and shivering were the most frequently mentioned as dengue symptoms (Table 3). In general, the responders showed knowledge about dengue as a disease, but few knew that the dengue virus has the same name as the disease (Table 3). Furthermore, most were aware that dengue is spread due to the mosquito bite (Table 3). Few knew that although the mosquito must be infected to transmit the disease, not all mosquitoes are infected (Table 3). Not many people specified that the *Aedes* mosquito is the mosquito involved in dengue virus transmission. The sources of acquiring the knowledge were mainly the television and the internet, highlighting the need for effective school programs to bring knowledge to the community (Table 4 provides the responder’s source of information about dengue and *Ae. aegypti* mosquitoes). Respondents declared having obtained knowledge regarding dengue and *Ae. aegypti* through television (159, 81.96%) internet (151, 77.84%) and school (64,32.99%).

**Table 3.**
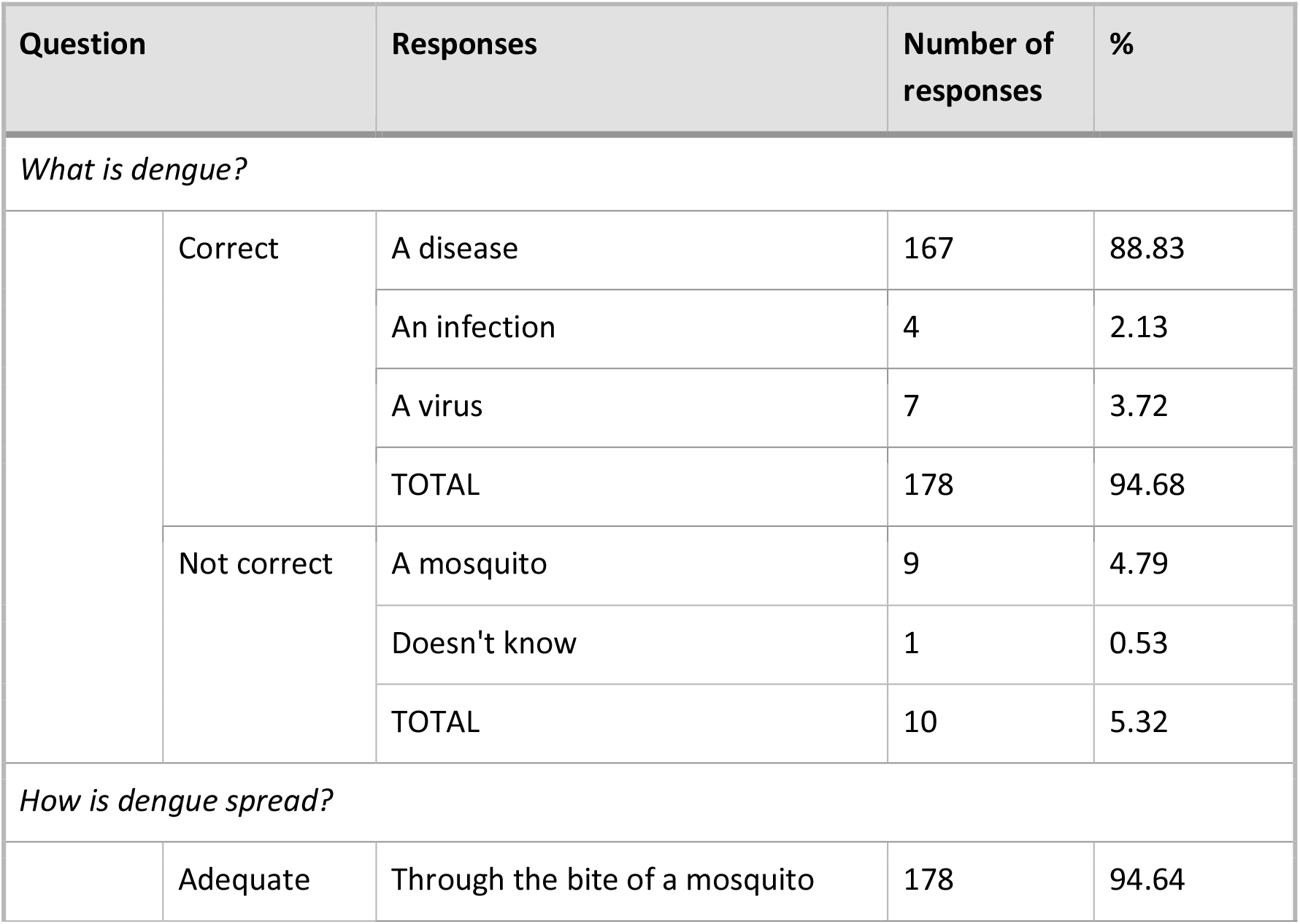
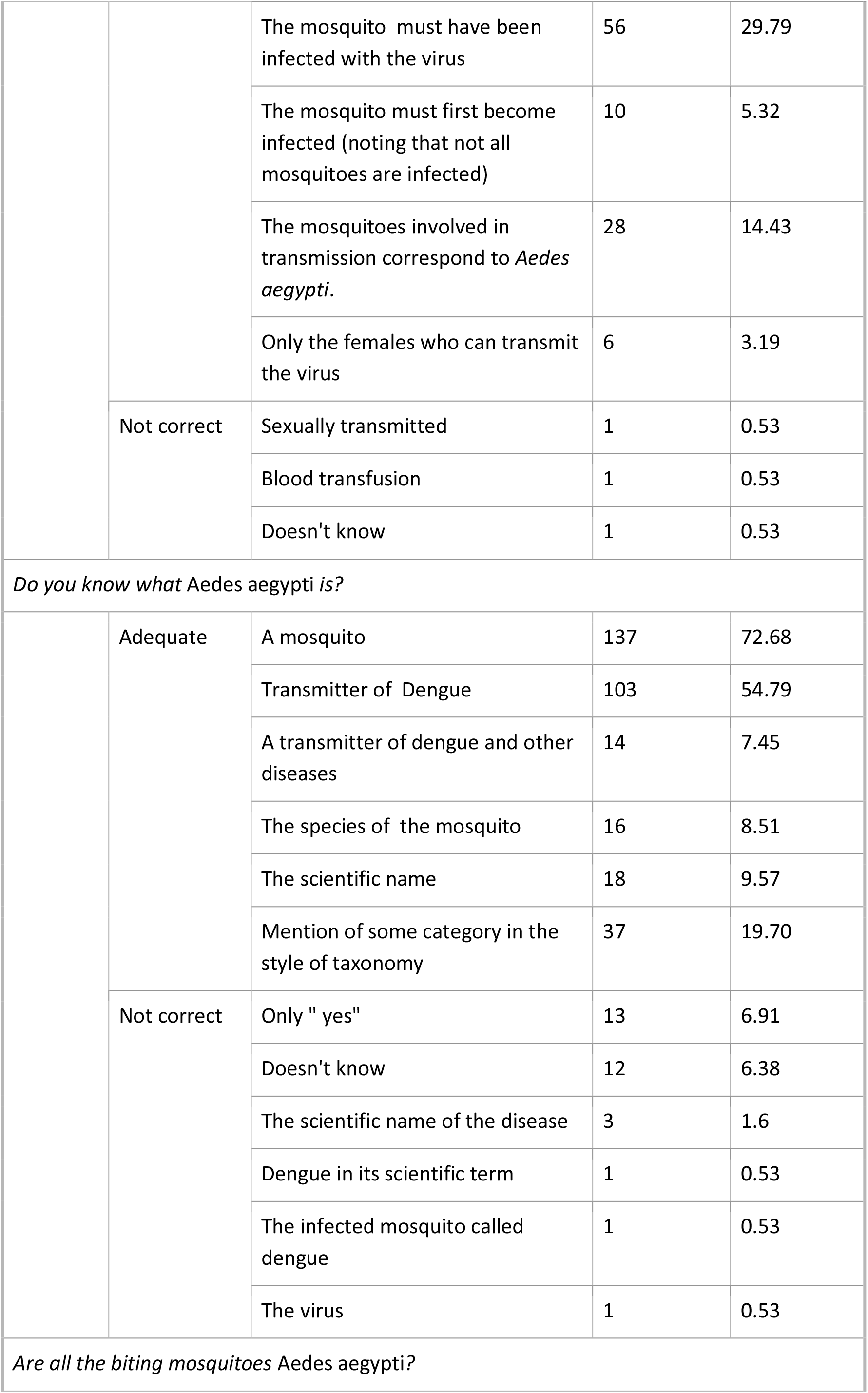
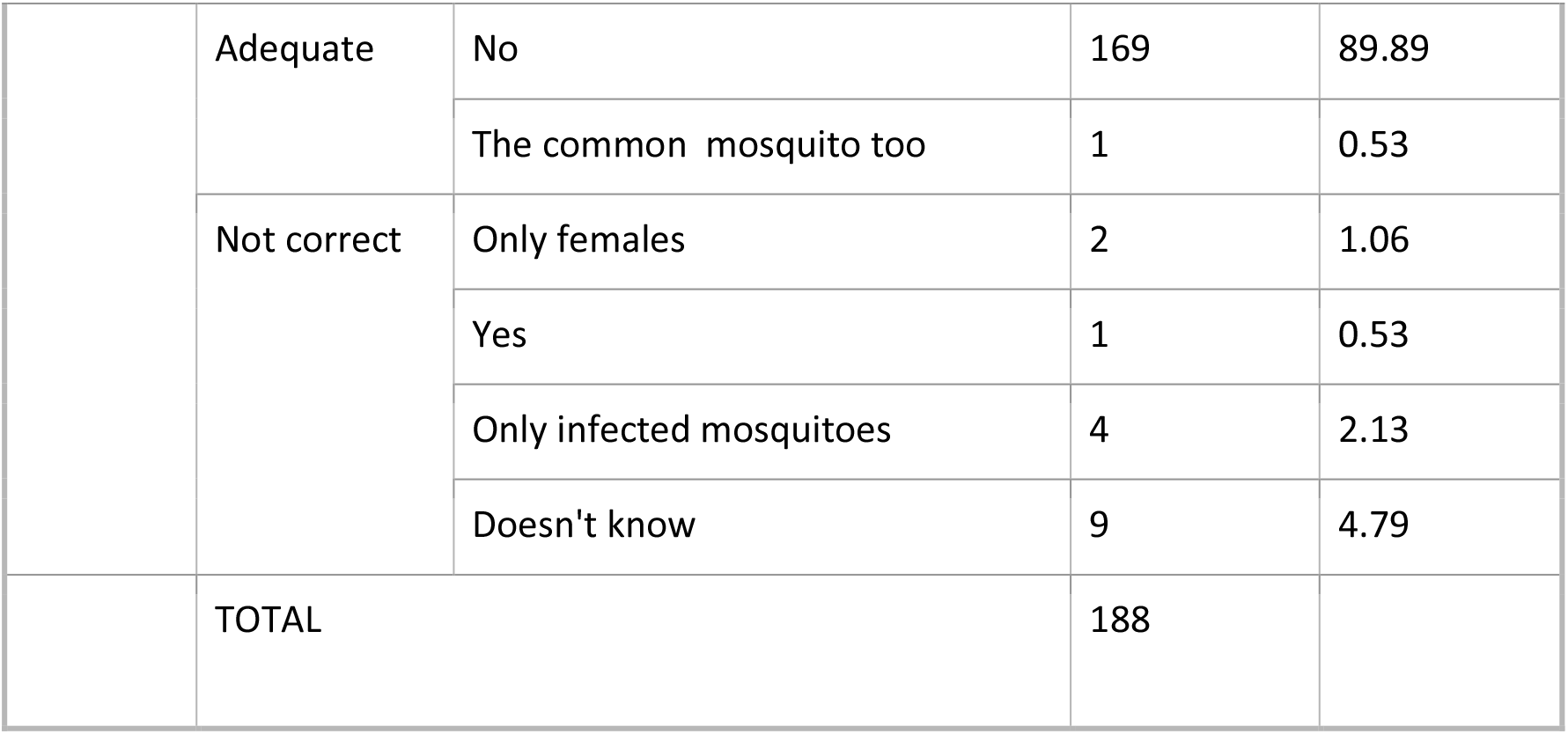
Information provided by respondents on vector-borne diseases knowledge.

| Question |  | Responses | Number of responses | % |
| --- | --- | --- | --- | --- |
| What is dengue? |  |  |  |  |
|  | Correct | A disease | 167 | 88.83 |
|  |  | An infection | 4 | 2.13 |
|  |  | A virus | 7 | 3.72 |
|  |  | TOTAL | 178 | 94.68 |
|  | Not correct | A mosquito | 9 | 4.79 |
|  |  | Doesn't know | 1 | 0.53 |
|  |  | TOTAL | 10 | 5.32 |
| How is dengue spread? |  |  |  |  |
|  | Adequate | Through the bite of a mosquito | 178 | 94.64 |
|  |  | The mosquito must have been infected with the virus | 56 | 29.79 |
|  |  | The mosquito must first become infected (noting that not all mosquitoes are infected) | 10 | 5.32 |
|  |  | The mosquitoes involved in transmission correspond to *Aedes aegypti*. | 28 | 14.43 |
|  |  | Only the females who can transmit the virus | 6 | 3.19 |
|  | Not correct | Sexually transmitted | 1 | 0.53 |
|  |  | Blood transfusion | 1 | 0.53 |
|  |  | Doesn't know | 1 | 0.53 |
| Do you know what *Aedes aegypti* is? |  |  |  |  |
|  | Adequate | A mosquito | 137 | 72.68 |
|  |  | Transmitter of Dengue | 103 | 54.79 |
|  |  | A transmitter of dengue and other diseases | 14 | 7.45 |
|  |  | The species of the mosquito | 16 | 8.51 |
|  |  | The scientific name | 18 | 9.57 |
|  |  | Mention of some category in the style of taxonomy | 37 | 19.70 |
|  | Not correct | Only " yes" | 13 | 6.91 |
|  |  | Doesn't know | 12 | 6.38 |
|  |  | The scientific name of the disease | 3 | 1.6 |
|  |  | Dengue in its scientific term | 1 | 0.53 |
|  |  | The infected mosquito called dengue | 1 | 0.53 |
|  |  | The virus | 1 | 0.53 |
| Are all the biting mosquitoes *Aedes aegypti*? |  |  |  |  |
|  | Adequate | No | 169 | 89.89 |
|  |  | The common mosquito too | 1 | 0.53 |
|  | Not correct | Only females | 2 | 1.06 |
|  |  | Yes | 1 | 0.53 |
|  |  | Only infected mosquitoes | 4 | 2.13 |
|  |  | Doesn't know | 9 | 4.79 |
|  | TOTAL |  | 188 |  |

**Table 4.** Information provided by respondents on How do they know about dengue and *Aedes aegypti*.

|  | TV | INTERNET | SCHOOL | OTHER | n | % |
| --- | --- | --- | --- | --- | --- | --- |
|  | * |  |  |  | 19 | 10.11 |
|  |  | * |  |  | 20 | 10.64 |
|  |  |  | * |  | 4 | 2.13 |
|  | * | * | * |  | 26 | 13.83 |
|  | * | * |  |  | 15 | 7.98 |
|  |  | * | * |  | 22 | 11.70 |
|  | * |  |  | * | 54 | 28.72 |
|  |  | * |  | * | 5 | 2.66 |
|  |  |  | * | * | 4 | 2.13 |
| n | 153 | 118 | 64 | 56 |  |  |
| % | 81.38 | 62.77 | 34.04 | 29.79 | 0 | 0 |

### Perception of dengue risk

Regarding perceptions of dengue, the majority (56%) affirm that dengue is a severe illness and 21% a moderate one. When people were asked how feasible they think it is to prevent dengue, most (59%) answered that it is easy while 38% said that it is relatively difficult. Two-thirds (67%) of respondents admitted that individuals play an important role in the prevention of dengue.

### Household control activities

Regarding mosquito control activities, 90% of respondents reported turning containers with water upside down and/ or eliminating them, 64% mow the grass, and 33% add chemicals to the pool. In addition, 92% claimed to use repellents or spirals to avoid mosquitoes, while 19% fumigate their homes. Regarding physical barriers, the majority (76%) acknowledged that they do not have window screens and 91% do not have door screens to prevent the entry of mosquitoes into their homes.

In addition to the aforementioned preventive measures, the respondents also reported organizing and cleaning the house and yard, disinfecting, and using insecticides. People also reported cleaning containers that contain water such as water bowls for pets, vases with aquatic plants, cleaning gutters, and checking that drains and garbage bags do not accumulate water.

### The environment surrounding the household

Regarding the environment surrounding households, abandoned urban land is often used to discard solid waste in urban neighborhoods. One-third (36%) of the surveyed households reported an informal dumping area near their dwelling. An additional 15% of the respondents had a formal garbage dump near their dwelling. We considered water channels and Suquia River as risk places because people throw trash along the shores, generating mosquito breeding sites; one-third (31%) of the respondents lived near water channels, and 21% lived near the Suquía River. Most people reported observing mosquitoes both inside and outside of the home (59%); however, some reported mosquitoes mostly outside their houses in the garden, backyard, or even open galleries (21%); and others (20%) reported mosquitoes only inside their dwelling. The majority (88%) reported seeing mosquitoes every day during the warm season in 2020, during the epidemic.

### Logistic regression analysis

In relation to knowledge about dengue, the majority (94.68%) responded corrected that dengue was either a disease (88.83%), an infection (2.13%) or a virus (3.72 %) Regarding how dengue is spread, 183 (94.33%) respondents indicated that it occurs through the bite of a mosquito. One-third (30.41%) indicated that the mosquito must have been previously infected with the virus, and 12 respondents (6.19%) pointed out that the mosquito must first become infected, noting that not all mosquitoes are infected. 28 (14.43%) respondents noted that the mosquitoes involved in transmission correspond to *Ae. aegypti*. Few people (3.19%) mentioned that it is only the female mosquitoes who can transmit the virus. When questioned whether the respondent knows what *Aedes aegypti* is, most (72.68%) answered correctly that it was a mosquito, and most (54.12%) mentioned that it was a transmitter of dengue. Several people (n=14) indicated that it also transmitted other diseases and others reported (n=16) that it was the species of the mosquito, or the scientific name (n=19). Most of the respondents knew that all the biting mosquitoes are not *Aedes aegypti*, (90.20%). A single respondent mentioned the existence of another mosquito, “the common mosquito”, which would correspond to *Culex quinquefasciatus*, which is very common in Córdoba.

Respondents correctly identified dengue symptoms such as fever, muscle pain, headache, etc. (Table 5). Taking into account the WHO characterization of symptoms, the variable ARS (Adequate answers about symptoms) was generated that is worth 1 when the respondent answered correctly and zero in another case. Based on the formulated hypothesis we expected to find that the correct knowledge of the symptoms of dengue disease depends on the educational level reached by each person. The association between ARS (Adequate responses about symptoms) and the EL (Educational level) of the responders (Table 6) showed that was partially correct. Although it was shown that at the graduate level there is a notable tendency to correctly indicate dengue symptoms, even more than expected (χ2= 21.63, p<0.001), there is no significative differences between the percentage of people that answer correctly and those who do not in the rest of the levels (Table 6).

**Table 5.**
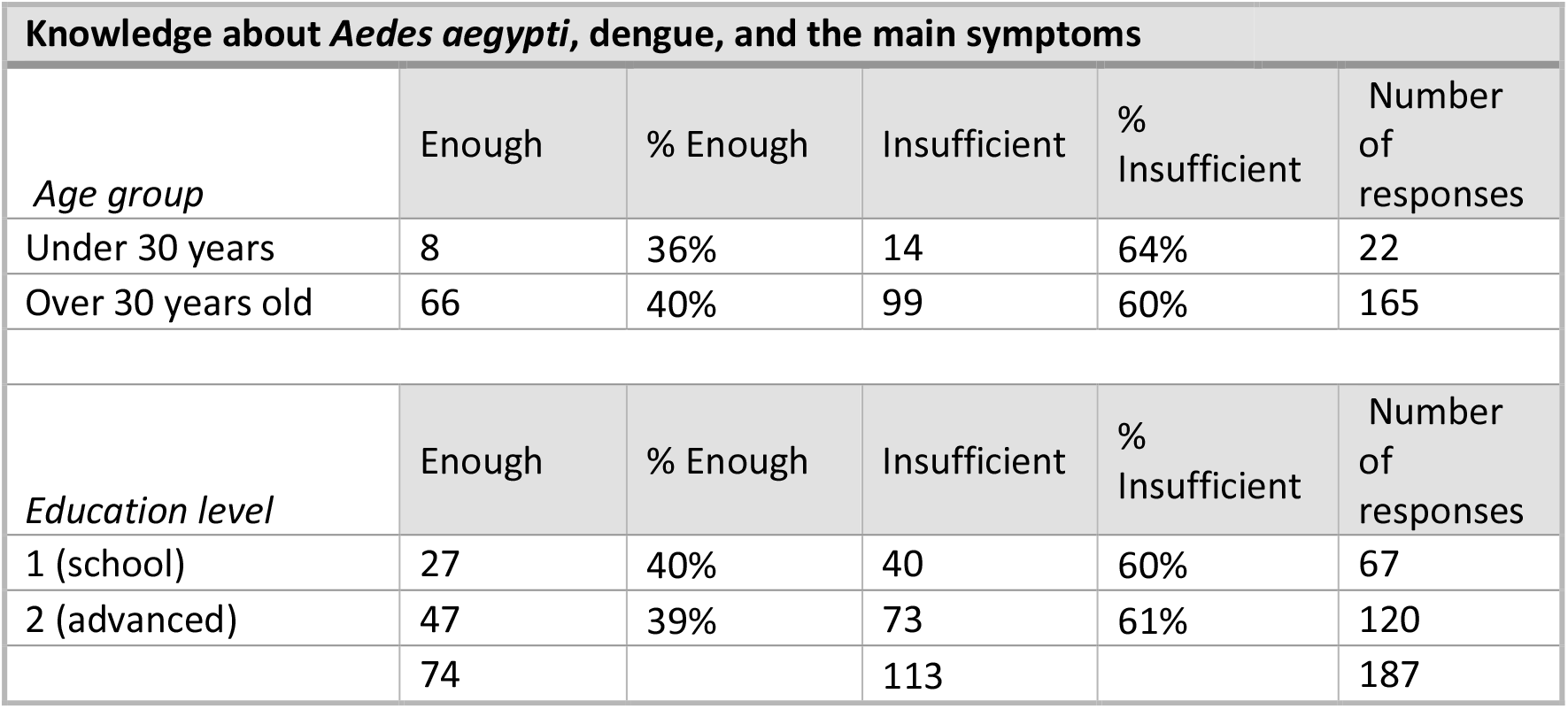
Knowledge about *Aedes aegypti*, dengue, and the main symptoms.

**Table 6.** Association between ARS (Adequate responses about symptoms (1: adequate; 0: not adequate) variables and the Educational level (EL) of the respondent.

| E L | ARS | n | % | $\chi^2$ | p-value |
| --- | --- | --- | --- | --- | --- |
| Primary | 0 | 2 | 1.15 | 0.53 | 0.4652 |
|  | 1 | 2 | 1.15 |  |  |
| Secondary | 0 | 27 | 15.52 | 4.68 | 0.0306 |
|  | 1 | 27 | 15.52 |  |  |
| Tertiary | 0 | 20 | 11.49 | 3.33 | 0.0679 |
|  | 1 | 20 | 11.49 |  |  |
| Graduate | 0 | 1 | 0.57 | 21.63 | <0.0001 |
|  | 1 | 68 | 39.08 |  |  |
| Postgraduate | 0 | 2 | 1.15 | 0.00 | >0.9999 |
|  | 1 | 5 | 2.87 |  |  |

n: number of respondents in each category; %: observed percentage of respondents belonging to each category; $\chi^2$ : comparison of observed and expected frequencies in each category.

When comparing knowledge about dengue and the vector (Table 7), and good preventive practices in relation to the educational level (EL) achieved and group age, the analysis indicated no significant differences. Regardless of this, we observed that a significant number of surveyors (84.5%) develop good preventive practices.

**Table 7.** Knowledge about *Aedes aegypti*, dengue, and the main symptoms.

| | | Enough | | Insufficient | | $\chi^2$ | p-value |
| --- | --- | --- | --- | --- | --- | --- | --- |
|  |  | n | % | n | % |  |  |
| Age group | 18–30 years | 8 | 36.4 | 14 | 63.6 | 0.11 | 0.7432 |
|  | >30 years | 66 | 40 | 99 | 60 |  |  |
| Educational level | Secondary | 27 | 40.3 | 40 | 59.7 | 0.02 | 0.8794 |
|  | Superior | 47 | 39.2 | 73 | 60.8 |  |  |
| Total |  | 74 | 39.6 | 113 | 60.4 | 4.12 | 0.0423 |

n: number of respondents in each category; %: observed percentage of respondents belonging to each category; $\chi^2$ : comparison of observed and expected frequencies in each category.

### About the Attitude Activity

With regards to the activity involving students’ attitude of exploring their backyards and/or their gardens and even inside their homes, searching for containers that hold water and could be potential mosquitoes breeding places, only 94 students (39 %) completed the activity and documented the activity with photography that was sent to their teachers. The students were able to identify mosquito breeding sites such as pots, buckets, bottles, drinkers, jars, vases, among others (Fig. 2). 89% of the students found water inside the containers and of these, 17% found the presence of mosquito larvae, although it was impossible to know if they were *Ae. aegypti*.

**Fig. 2.**
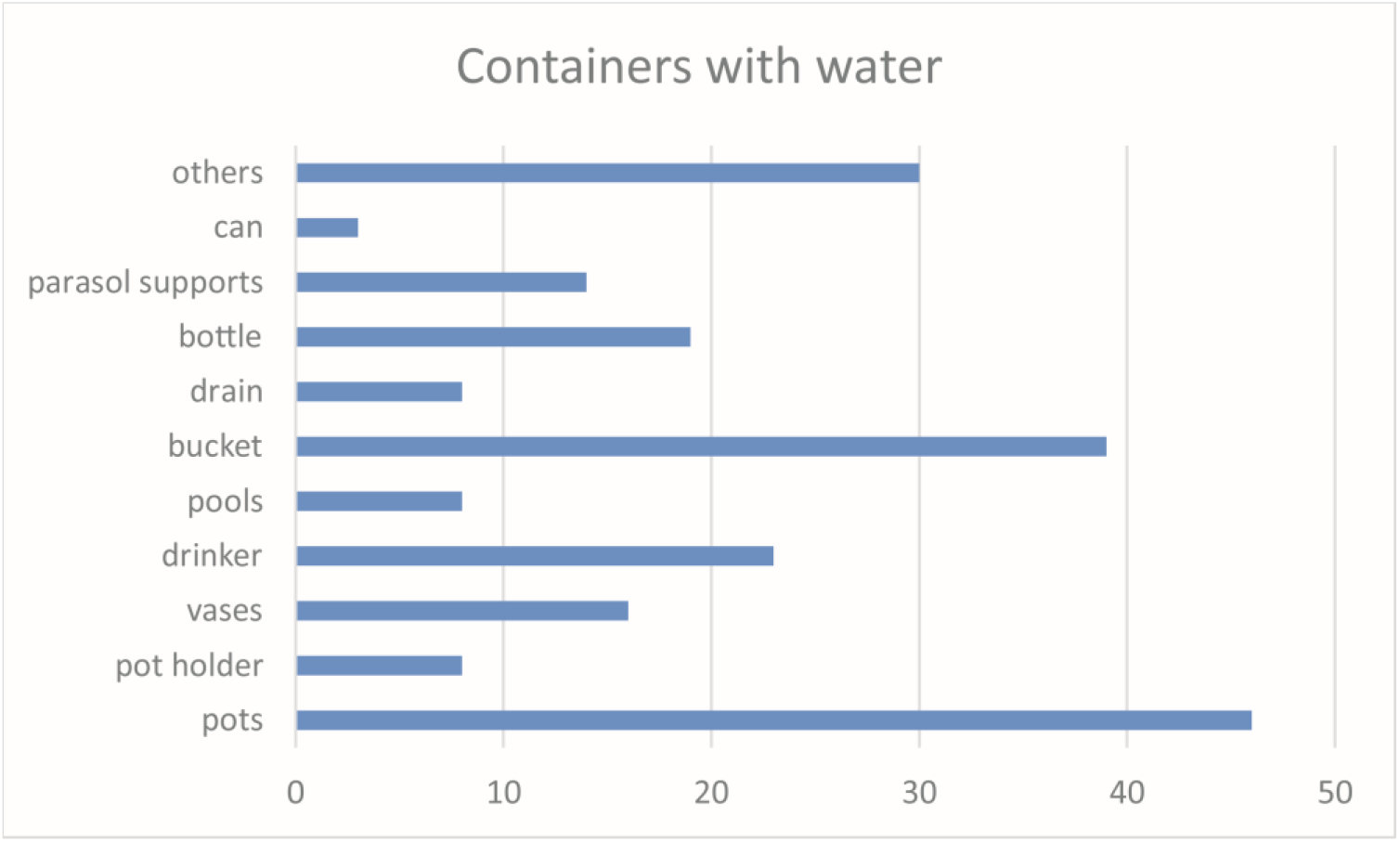
Potential breeding sites. Number and type of containers with water found in the dwellings of the students who carried out the activity.

### About the Action

Finally, students put into practice the knowledge acquired by eliminating containers as well as storing them indoors without water, documenting with photos that were sent to the teacher in each of the schools involved. 61 students (65%) gave their opinions about the whole activity describing a positive impact on their family environment and future decisions with respect to vector management.

## Discussion

Considering that community awareness and education are critical factors for *Ae. aegypti* mosquito’s management, we thought of this pilot online experience during the COVID-19 pandemic quarantine. Student-led science engaged the teachers and involved students during online classes and the biggest dengue outbreak in our region’s history. As we already know, dengue prevention depends on effective vector management. According to the World Health Organization [28], sustained community involvement can improve vector management efforts substantially. Although, mosquito educational interventions by the government in temperate areas of Argentina are done just during adult vector activity months when adult mosquitoes batter the people (November to April mainly), they are not sustained all year long as could be in tropical areas where dengue risk is all the yearlong.

As expressed in the surveys, people seem to have good knowledge about the vector and the disease in Córdoba city. A striking case is that observed in Malaysia where despite being in constant contact with the disease, since the risk of outbreaks is frequent or continuous [29], research shows that there is insufficient knowledge about the cause of the disease and how it is transmitted, as well as about the activity of the vector and the breeding sites where it develops [30]. Our results revealed the knowledge of people about *Ae. aegypti* mosquito but the lack of knowledge about *Ae. aegypti* as a vector that could be infected with the dengue virus. The need for appropriate knowledge transference to the community focuses on females like those who bite the humans and transfer the virus if they are infected. In accordance with McNaughton et al. [31], controlling dengue fever relies heavily upon the actions of residents as well as community education and awareness of the risks, if people do not know the cause of the disease and how it is transmitted later; therefore, appropriate practices cannot be implemented to prevent it. The mosquito vector seems to be known better by television and less by the school, indicating that television is an essential source of knowledge, as other studies assert [30, 32]. This generates an impact on the audience. It makes us think of television as a critical factor in vector-borne corrected knowledge transmission, awareness and spreading information by internet.

In addition, these could be highlighting the need of schools to add in their programs as main contents about mosquito biology and vector-borne diseases to raise awareness of a right knowledge diffusion to families and hence the community aiming to prevent vector breeding sites as well as the disease in a sustainable time as suggested by WHO. In fact, in accordance with previous studies, which evidenced the role of education leading behavior changes and increasing community knowledge after vector-borne focus education [11, 12]. About dengue perception, in Córdoba, the surveyors recognize the illness’s importance and severity. These findings highlight a considerable difference concerning other communities in other countries like Colombia, which think it is not an illness but just regular flu (mild flu) or common cold [33]. Likewise, in a study made in Malaysia, most survey respondents recognized the common symptoms or signs of the disease but not the complications they may manifest [30]. In tropical areas, people are used to the large number of dengue cases that occur each year because of being in an area where the presence of the vector and the virus circulation is constantly unlike temperate Córdoba city, with vector activity during the warmer months. For this reason, people could underestimate its importance.

Our study highlights people understanding of dengue prevention as a responsibility of each individual, including schools and government, unlike other studies in which most people place responsibility on the government and health care authorities besides each household [32]. In fact, in our study, people affirm that school has a fundamental role in raising awareness and educating. For its part, the government communicates, gives information, fumigates, and takes control of the vector situation. With this, phrases like ^..^joint work^..^ and ^..^social responsibility^..^ should be pointed out.

People movement is one of the main causes of disease expansion [34]; we reported that people with travel, most travel recently to neighboring countries like Brazil, that presents more than 2,226,900 dengue cases during 2019 with the highest impact in history [5]. This situation increases the risk of income imported cases to the country.

Indeed, areas of southern Brazil reported co-circulation of dengue virus 1, 2, and 4 until epidemiological week 23 [35].

It is important to mention that all the prevention methods that people refer to are recommendations from the media highlighting their relevance to people’s awareness.

## Conclusion

Having good knowledge alone is not sufficient for dengue control while perception towards barriers, self-efficacy, and cues to act among respondents can be further strengthened. As dengue is related to health behavior, future prevention and control programs can be designed to encourage communities to make decisions and behavior changes for dengue and dengue prevention effectively. Through this study, we encourage the need of adding school programs to engage the community and transmit the constant implications of the mosquito-borne disease, and the need for vector mosquito management together as a community. We encourage the need for schools and government campaigns together, with the engagement of journalists and television as the means for people to get knowledge on mosquito *Ae. aegypti* and dengue diseases massively. We need to inculcate positive attitudes on the population and cultivate better preventive practices to eliminate the breeding places for mosquitoes’ vectors and therefore diminished the possibility of dengue fever circulating in our city.

## Supplementary information

**Additional file 1.** Survey instrument including questions regarding the identification of vector-borne diseases, socio-ecological factors, knowledge, and perception of dengue, household vector prevention and control actions, as well as the household environmental surrounding. (Additional file 1.docx)

**Additional file 2.** Additional details about training of students to learn about the vector and carry out the Attitude activity. (Additional file 2.docx)

## Declarations

**Ethics approval and consent to participate:** Not applicable.

**Consent for publication:** Not applicable.

**Availability of data and materials:** Not Applicable

## Author Contributions

**Elizabet L. Estallo:** Conceptualization, Methodology, Investigation, Formal Analysis, Writing - original draft, Writing - review & editing. **Magali Madelon:** Investigation, Formal Analysis, Writing - original draft, Writing - review & editing. **Elisabet M. Benitez:** Investigation, Formal Analysis, Writing - original draft, Writing - review & editing. **Anna M. Stewart-Ibarra:** Writing - original draft, Writing - review & editing. **Francisco F. Ludueña-Almeida:** Methodology, Investigation, Formal Analysis, Writing - original draft, Writing - review & editing.

## Acknowledgments

The authors wish to acknowledge the authorities and teachers from four high school “Jóvenes Argentinos”, “Centro Educativo Nuevo Siglo”, “Instituto Nuestra Señora de Fatima” and “Instituto Jesús María” for their engagement and collaboration doing Citizen science through this study. This research was supported by the Science and Technology Ministry of Córdoba Province (MINCYT) grant GRFT 2019/000298 and the Secretary of Science and Technology (SECyT-UNC) grant 33620180100735CB. Also, we thank Daniela Tinunin and Agustina Ortiz for their help in survey development and part of the analysis. Daniela Tinunin developed her undergraduate thesis to develop as a biologist with the direction of Dra. Estallo and Dr. Ludueña- Almeida. Her colaboration is part of an intership carried out in the Centro de Investigaciones Entomológicas de Córdoba, Facultad de Ciencias Exactas, Físicas y Naturales, Universidad Nacional de Córdoba. ELE is a member of the Consejo Nacional de Investigaciones Científicas y Técnicas (CONICET) from Argentina and EMB is a PhD Student with scholarship support from CONICET.

